# Aluminium alters excitability by inhibiting calcium, sodium and potassium currents in bovine chromaffin cells

**DOI:** 10.1101/2023.01.25.525351

**Authors:** Andrés M. Baraibar, Ricardo de Pascual, Victoria Jiménez Carretero, Natalia Hernández Juárez, Itxaso Edurne Aguirregabiria Alonso, Jesús M. Hernández-Guijo

## Abstract

Aluminium (Al^3+^) has long been related to neurotoxicity and neurological diseases. This study aims to describe the specific actions of this metal on cellular excitability and neurotransmitter release. Al^3+^ reduced intracellular calcium concentrations around 25% and decreased catecholamine secretion in a dose-dependent manner, with an IC_50_ of 89.1 μM. Al^3+^ blocked calcium currents in a time- and concentration-dependent manner with an IC_50_ of 560 μM. This blockade was irreversible, since it did not recover after wash-out. Moreover, Al^3+^ produced a bigger blockade on N-, P- and Q-type calcium channels subtypes (69.5%) than on L-type channels subtypes (50.5%). Sodium currents were also inhibited by Al^3+^ in a time- and concentration-dependent manner, 24.3% blockade at the closest concentration to the IC_50_ (419 μM). This inhibition was reversible. Voltage-dependent potassium currents were non-significantly affected by Al^3+^. Nonetheless, calcium/voltage-dependent potassium currents were inhibited in a concentration-dependent manner, with an IC_50_ of 447 μM. This inhibition was related to the depression of calcium influx through voltage-dependent calcium channels subtypes coupled to BK channels. In summary, the blockade of these ionic conductances altered cellular excitability that reduced the action potentials firing and so, the neurotransmitter release and the synaptic transmission. These findings prove that aluminium has neurotoxic properties because it alters neuronal excitability by inhibiting the sodium currents responsible for the generation and propagation of impulse nerve, the potassium current responsible for the termination of action potentials, and the calcium current responsible for the neurotransmitters release.

## 1. Introduction

Aluminum (Al^3+^) is an extremely ubiquitous element, being the third most abundant on the Earth’s crust (Krewski et al., 2009). Although this metal has no physiological role in any metabolic processes, humans are constantly exposed to it. Al^3+^content in water results from a combination of factors including, mobilization, the weathering of rocks and minerals (Campbell et al., 1992), and the use of aluminium minerals in water purification (Krewski et al., 2009, Krypinska, 2020). Its presence in foods may be either natural or attributable to the use of cookware, utensils and wrappings containing Al^3+^, in addition to additives (Piepponen, 1992; Sepe et al., 2001). Drugs (phosphate binders, antacids, buffered analgesics, antidiarrheal and antiulcer drugs) represent another important source of Al^3+^ intake (Krewski et al., 2009, Crisponi et al., 2013). Likewise, pesticides, disinfectants, cosmetics and anti-perspirants are sources of Al^3+^ exposure (Soni et al., 2001; Saived and Yokel, 2005).

Estimates of Al^3+^ intake range from 0.7 mg/day for 6-to 11-month-old children to 11.5 mg/day for 14-to 16-year-old males. For adult men and women these estimates are 8-9 and 7 mg/day (Soni et al., 2001). For the general population, intake from food (10^−4^ mg/kg body weight (b.w.)/day) dominates that from drinking water (1.7 × 10^−5^ mg/kg b.w./day) and inhalation exposure (6.9 × 10^−6^ mg/kg b.w./day) (1). Besides, other sources, for example, antiacid treatment and buffered aspirin, amount to 3.1 × 10^−1^ and 4.3 × 10^−2^ mg/kg b.w./day, respectively (Soni et al., 2001). In conclusion, the average consumption of the metal could be around 25 mg/day (Soni et al., 2001). Nevertheless, many products act as Al^3+^ accumulators (Soni et al., 2001) and provide more Al^3+^ when consumed in one meal as part of the standard daily diet. Taking into consideration that some products are also often consumed, this leads to the potential for even higher Al^3+^ daily intake. Examples of products that contain high amounts of Al^3+^ are frozen pizzas, baked goods (Saiyed and Yokel, 2005), chocolate or tea infusions and soft drinks (Sepe et al., 2001). The Joint FAO/WHO Expert Committee on Food Additives (JEFCA) established the tolerable weekly intake for aluminium in 2 mg/kg of b.w./week. Notwithstanding, in many countries, considerable amounts of people, especially children, surpass this limit, being at risk of Al^3+^ toxicity (Crisponi et al., 2013; Fekete et al., 2013; Yang et al, 2014). In addition, an important group of vulnerable people are workers in occupations related to scrap metal recycling, mineral extraction and processing, the use of compounds and products containing Al^3+^; cutting and welding of Al^3+^ (Igbokwe et al., 2020; Valkonen and Aitio, 1997), whose exposure levels are significantly higher (6 × 10^−3^ mg/kg b.w./day) (Klotz et al., 2017). This occupational exposure has been related to neurological problems, like loss of coordination, memory and balance (Krewski et al., 2009).

It exists a clear association between high levels of Al^3+^ in the CNS and neurotoxicity (Krewski et al., 2009; Kawahara and Kato-Negishi, 2011; Singla and Dhawan, 2012). Al^3+^ is absorbed (Jaishankar et al., 2014), it reaches systemic blood circulation where is transported bound to transferrin (Nagaoka and Maitani, 2005), the complex has the ability to cross the blood brain barrier (BBB) (Deloncle et al., 1990), and it distributes and accumulating mainly in bone, kidney and brain (Krewski et al., 2009; Nebeker and Coburn, 1986). This event alone may directly alter the BBB function, allowing the passage of many compounds (Deloncle et al., 1990). Additionally, the transferrin receptor in the brain enables the entrance of iron, so, Al^3+^ interferes with normal iron homeostasis, disrupting iron-dependant cellular processes in the CNS (Roskams and Connor, 1990). Notwithstanding, the main mechanism of Al^3+^ neurotoxicity is the reactive oxygen species production, giving rise to oxidative stress (Sánchez-Iglesias et al., 2009) and mitochondrial dysfunction (Kumar and Gill, 2014). It has been reported a relation between chronic high-Al^3+^ diet and neuronal loss (Kihira et al., 2002); and many studies have related Al^3+^ as a risk factor in Alzheimer’s disease (Walton and Wang, 2009; Exley et al., 1993, Mantyh et al., 1993; Savory et al., 1995; Bouras et al., 1992).

The main symptoms of Al^3+^ toxicity are digestive (nausea, vomiting and diarrhea), motor (weakness, fatigue, motor disturbances, osteomalacia) and neurological (depression, dementia, hallucinations, decreased intellectual function, inability to concentrate, speech impairment and language, and epileptic seizures) (Soni et al., 2001; Jaishankar et al., 2014). Al^3+^ was first considered toxic after the discovery of high levels of this metal in brain tissue samples from encephalopathy patients (Alfrey, 1993) who were exposed to Al^3+^ during dialysis (Parkinson et al., 1981). Since then, Al^3+^ has been linked to diseases such as Alzheimer’s disease (Chin-Chan et al., 2015; Mcdermott et al., 1979), Parkinson’s disease (Uversky et al., 2001) and multiple sclerosis (Mold et al., 2018) among others (Linhart et al., 2017; Li et al., 2011; Novaes et al., 2018; Zhu et al., 2013; Mailloux et al., 2011; Martinez et al., 2017). In fact, it has been reported higher incidence of Alzheimer’s disease in regions with high levels of Al^3+^ in drinking water (Flaten, 2001; Forbes and McLachlan, 1996), suggesting an important role of this metal in the pathogenesis of the disease. Majority of reports regarding Al^3+^ toxicity is related to chronic events, acute Al^3+^ intoxication is also very important because of the incidence of fatal cases (Soni et al., 2001).

Even low levels of Al^3+^ exposure can increase its concentration in the CNS (Martinez et al., 2017), triggering neurotoxicity and degenerative brain disease, resulting in cognitive damage and promoting the onset and progression of neurodegenerative diseases (McLachlan et al., 1986; Fernandes et al., 2020) as the Alzheimer’s disease (AD) (Flaten et al., 2001). Al^3+^ increases amyloid-β production by increasing amyloid precursor protein levels (Lin et al., 2008), reduces amyloid-β degradation by inhibiting the cathepsin B (Sakamoto et al., 2006), and produces a conformational change in amyloid-β enhancing its aggregation (Kawahara et al., 2001). Al^3+^ levels are increased in patients with Alzheimer’s disease, existing a co-localization with the amyloid-β plaques in the human brain (Crapper et al., 1973; Mcdermott et al., 1979).

To date, little data is available on how Al^3+^ affects cell excitability and synaptic transmission. Low levels of Al^3+^ have been reported to inhibit calcium currents (Busselberg et al., 2009; Platt and Busselberg, 1994), reducing the amplitude of excitatory postsynaptic potentials (EPSP) (Platt et al., 1995); while higher levels improve calcium currents (Chen et al., 2005), increasing the amplitude of the EPSPs (Platt et al., 1995). While its effects on other ionic currents, such as sodium and potassium currents, seem to be scarce (Meiri and Shimoni, 1991). This metal has been reported to alter long-term potentiation (LTP) (Song et al., 2014), the neurophysiology basis of learning and memory (Wang et al., 2010; Cao et al., 2016); processes where Ca^2+^/calmodulin-dependent protein kinase II and cAMP-dependent protein kinases are important (Harris et al., 1994; Frey et al., 1993; Malenka et al., 1989, Matthis et al., 1993). It has been described that Al^3+^ interferes with the phosphorylation processes that affect cAMP (Johnson and Jope, 1987), alters Ca^2+^/calmodulin-dependent enzymes (Levine et al., 1990) and interferes with G proteins (Haug et al., 1994). Furthermore, Al^3+^ alter calcium homeostasis (Sautu et al., 2018); so, the elevation of the calcium concentration, as a result of the release from cellular organelles or calcium-binding proteins, in combination with the altered absorption of calcium due to the interaction between Al^3+^ and Mg^2+^-dependent ATPase (Suhayda and Haug, 1984; Siegel and Haug, 1983), may modify the release of neurotransmitters. This fact, together with the inhibition of the enzyme acetylcholinesterase exerted by aluminium (Zatta et al., 2002) leads to modulate cholinergic neurotransmission (Szutowicz, 2001, Yellamma et al., 2010).

Chromaffin cells, as sympathetic neurons are developed from the neural crest. They are excitable cells with neuron-like electrical properties (Biales et al., 1976) with the capacity to synthesize, store and release adrenaline and noradrenaline (for review, see Huber, 2006). They are one of the most popular and widely used cellular models for investigating the molecular mechanisms underlying cellular excitability and neurotransmitter release (Tischler, 2002).

Based on the data that daily intake of Al^3+^, which leads to the possibility of easily exceeding the admitted toxic levels, and the ability of this metal to trigger neurotoxicity and its implication in pathologies that affect the central nervous system; in combination with previous data indicating a possible role of Al^3+^ as a modulator of synaptic transmission; the present work, using chromaffin cells, aims to research how the alterations of cellular excitability and synaptic transmission induced by Al^3+^ underlies their effects reported on central nervous system, through the analysis of the effects of Al^3+^ on the main ionic currents responsible for the generation, transmission and completion of action potentials firing, as well as neurotransmitter release. Here, we demonstrate that Al^3+^ reduced intracellular calcium and decreased catecholamine secretion; and blocked calcium currents in a time-irreversible- and concentration-dependent manner; sodium currents were also reversible-inhibited; voltage-dependent potassium currents were not affected whereas calcium/voltage-dependent potassium currents were inhibited. This effects of ionic conductances drastically modifies cellular excitability.

## 2. Materials and methods

### 2.1 Isolation and culture of bovine chromaffin cells

The care and use of animals were carried out in accordance with the National Council on Animal Care and the European Communities Council Directive (86/609/ECC); and were approved by the local Animal Care Committee of Universidad Autónoma de Madrid (animal facilities ES280790000092).

Adrenal glands were obtained from a local slaughterhouse under the supervision of the local veterinary service. Bovine chromaffin cells were isolated by digestion of the adrenal medulla with collagenase (de Pascual et al., 2016). Briefly, cells were suspended in Dulbecco’s modified Eagle’s medium (DMEM), and were further supplemented with 5% foetal bovine serum, 50 IU/ml penicillin, in combination with 50 μg/ml streptomycin. Proliferation inhibitors (10 μM cytosine arabinoside, 10 μM fluorodeoxyuridine and 10 μM leucine methyl ester) were added to the medium to prevent excessive fibroblast growth. Cells were plated on 10 cm diameter Petri dishes 5 × 10^6^ cells in 10 ml of DMEM in order to carry out amperometry detection for secretion experiments. For patch-clamp studies, cells were plated on 1cm-diameter glass coverslips at low density (5 × 10^4^ cells per coverslip). For intracellular calcium measurements, cells were plated at a density of 2 × 10^5^ cell per well into 96-well plates. Cultures were maintained in an incubator at 37ºC in a water-saturated environment with 5% CO_2_. Cells were used 1-4 days after plating.

### 2.2 On-line measurement of neurotransmitters release

This experimental electrochemical method to measure exocytosis is based on the capacity of different substances to go through oxidation or reduction (Leszczyszyn et al., 1990).

Bovine chromaffin cells were scraped off carefully from the bottom of the Petri dishes with a rubber spatula and centrifuged at 120 g for 10 minutes. The cell pellet was then re-suspended in 200 μl Krebs-HEPES solution at 7.4 pH, that contained (in mM): 144 NaCl, 5.9 KCl, 1.2 MgCl_2_, 11 glucose, 10 HEPES, 2 CaCl_2_. Following, cells were placed in a microchamber (100 μl volume) for their cell perfusion at 37 ºC with Krebs-HEPES solution at 2 ml min^−1^. Under these conditions, the cell perfusion fluid emanating from the microchamber reached an electrochemical detector (model CH-9100; Metrohm AG, Herisau, Switzerland), equipped with a carbon fiber microelectrode, and placed just at the outlet of the microchamber. The catecholamines were oxidized at +0.65 V and the oxidation current was recorded with a frequency of 2 Hz, to monitor the amount of total catecholamine secreted. To stimulate the catecholamine secretion, a Krebs–Hepes solution containing KCl at 35 mM with isosmotic reduction of NaCl (35K^+^ solution), was applied in 5 s pulses at 1 min regular intervals at 37 ºC. After each stimulation, we measured in real time the catecholamine release by amperometry (Green and Perlman, 1981). Representative records of some experiments shown in this report were accomplished by importing the data obtained in ASCII format to the Origin 8.6 (Microcal) program. The total charge (Q) can be calculated by integrating the current versus time, using the Faraday’s law, expressed as Q = nNF, being n the number of electrons transferred in the redox reaction (n=2 for catecholamines), N the number of neurotransmitter molecules detected, and F the Faraday constant. Solutions were rapidly exchanged through electron-valves operated by a computer.

### 2.3 Electrophysiological recordings

Voltage-clamp and current clamp recordings were obtained with the whole-cell configuration of the patch clamp technique (Hamill et al., 1981). Recordings were made using patch pipettes of thin firepolished borosilicate glass (Kimax 51, Witz Scientific, Holland, OH), in order to obtain a final series resistance of 2−5 MΩ when filled with the standard intracellular solutions and mounted on the headstage of an EPC-10 patch-clamp amplifier (HEKA Electronic, Lambrecht/Pfalz, Germany), allowing cancellation of capacitative transients and compensation of series resistance. Data were acquired with a sampling frequency ranging between 5-10 kHz and filtered at 1-2 kHz. Recording traces with leak currents >100 pA or series resistance 20 MΩ were discarded. The P/4 protocol was used to discard linear leak and capacitive components. Data acquisition and analysis was performed using PULSE software (HEKA Electronic, Lambrecht/Pfalz, Germany).

Coverslips containing 5 × 10^4^ cells were placed on a chamber mounted on the stage of a Nikon Eclipse T2000 inverted microscope. During the seal formation with the patch pipette, the chamber was continuously perfused with a bubbled control Tyrode solution containing (in mM): 137 NaCl, 5 KCl, 1 MgCl_2_, 2 CaCl_2_, 10 HEPES/NaOH, (pH 7.4). Once the patch membrane was perforated and the whole-cell configuration of the patch-clamp technique was established, the cell was locally, rapidly and constantly superfused with an extracellular solution of similar composition to the chamber solution, but containing nominally 0 mM Ca^2+^ (to measure I_Na_), 10 mM Ca^2+^ (to measure I_Ca_) and 2 mM Ca^2+^ (to measure I_K_) (see Results for specific experimental protocols). For I_Na_ and I_Ca_ recordings, cells were dialyzed with an intracellular solution containing (in mM): 10 NaCl, 100 CsCl, 14 EGTA, 20 TEA.Cl, 5 Mg-ATP, 0.3 Na-GTP, 20 HEPES/CsOH (pH 7.3). To register I_K_, cells were internally dialyzed with that internal solution where CsCl and TEA were replaced by KCl. 5 μM tetrodotoxin (TTX) was added to external solution to measure I_Ca_; and TTX + 200μM Cd^2+^ to measure I_K._

To establish the perforated patch configuration, we used a pipette solution containing 50-100 ng/ml amphotericin B as permeabilizing agent (Calbiochem, Madrid, Spain) and a pipette-filling solution containing (in mM): 135 KCl, 4 NaCl, 1 MgCl_2_, 2 CaCl_2_, 5 EGTA, 20 HEPES/NaOH (pH 7.3). Amphotericin B was dissolved in dimethyl sulfoxide (DMSO, Sigma) and stored at -20 °C in stock aliquots of 50 μg/ml. Fresh pipette solution was prepared every 2 h. To facilitate the sealing, the pipette was first dipped in a beaker containing the internal solution and then back-filled with the same solution containing amphotericin B. Recording started when the access resistance decreased below 20 MΩ, which usually happened within 10 min after sealing (Rae et al., 1991). Series resistance was compensated by 80% and monitored throughout the experiment. Current-clamp experiments were analysed using Clampfit software (Molecular Devices, CA, USA). We have selected the next detection parameter value that the events must meet to be considered an action potential: time before a peak for baseline 20 ms (pretrigger) and period to search a decay time 25 ms (postrigger). Any action potential that did not have a clear morphology was discarded.

External solutions were rapidly exchanged using electronically driven miniature solenoid valves (The Lee Company, Westbrook, CO, USA) coupled to a multi-barrel concentration-clamp device, the common outlet of which was placed within 100 μM of the cell to be patched. The flow rate was 1 ml/min. All the experiments were performed at room temperature (22–24 °C). Only one chromaffin cell was used for a single experiment.

### 2.4 Measurements of [Ca^2+^]_c_ with Fluo-4AM

These experiments were carried out by using the fluorescent probe Fluo-4 AM (Thermo Fisher Scientific) and a microplate reader Fluostar Optima (BMG Labtech, Offenburg, Germany). After removing the medium, cells were incubated with the Ca^2+^ fluorescent probe Fluo-4 (Gibco-Invitrogen) (solution containing (in mM): 5.9 KCl, 144 NaCl, 1.2 MgCl_2_, 11 glucose, 10 HEPES/NaOH (pH 7.4) where 10 μM fluo-4-AM and 0.2% pluronic acid were included) for 45 minutes at 37ºC in dark. After this incubation period, cells were washed twice in dark, with the Krebs-HEPES buffer at room temperature. Fluorescence measurements were carried out using an excitation wavelength of 488 nm and recording the emission at 522 nm. AT the end of the experiment, cells were 10 min incubated with Triton X-100 (5%) to determine the maximum fluorescence (Fmax) and the incubated in the presence of MnCl_2_ (2 M, 10 min) to measure the minimum fluorescence (Fmin). Changes in [Ca^2+^]_c_ were calculated as percentage of the total fluorescence; Fx = (F_measured_-F_basal_) / (F_max_-F_min_) x 100. All experiments were performed at room temperature on cell from 1 to 3 days after culture.

### 2.5 Chemicals

AlCl_3_ was obtained from Sigma (Madrid, Spain). Collagenase type I was from Roche laboratory (Madrid, Spain), while DMEM, fraction V foetal bovine albumin and antibiotics were from Gibco (Madrid, Spain). Fluo-4-AM was obtained from Molecular Probes (Life Technologies), and the rest of the chemical reagents and solutions were from Sigma, Merck and Panreac Chemical. Nifedipine (Sigma) was prepared in stock solutions in ethanol (DMSO) at concentrations of 10 mM and protected from light. Toxins ω-conotoxin-GVIA (ω-CTx-GVIA) and ω-agatoxin-IVA (ω-Aga-IVA) (Peptide Institute, Osaka, Japan) were dissolved in distilled water at 0.1 mM stock concentration and stored frozen.

### 2.6 Statistical analysis

Data were expressed as means ± S.E.M. of the number of experiment done (n) from at least three different cell cultures. Student’s *t*-test or one-way ANOVA followed by Newman-Keuls multiple comparison test were used to determine statistical significance between means. For APs, analysis data were first subjected to a normality test (D’Agostino and Pearson Omnibus Normality test) and we found that some parameters followed a normal distribution and others did not. Thus, we applied a Mann-Whitney test in all cases. The statistical significance was established at *p* values smaller than 0.05 (*), 0.01 (**) and 0.001 (***). All analyses were performed using GraphPad Prism 6.01 software package.

## 3. Results

### 3.1. Al^3+^ decreases neurotransmitter release

In the experiment shown in Fig.1 cells superfused with Krebs-HEPES solution had a basal steady-stable spontaneous catecholamine release of about 20 nA. Upon challenge with solution containing 35 mM K^+^ (35K) administered in 10 s pulses, in order to trigger neurotransmitter secretion after direct depolarization of the cell membrane, secretion peaks of around 800 nA were obtained; these peaks show a constant neurotransmitter secretion when the K^+^ pulses were repeated at 5 min intervals (see control panel A). In these experiments (see panels B-D), the superfusion of the cells with a single concentration of Al^3+^ (30, 100, 300 and 1000 μM) from the K^+^ pulses P5 to P8 show a sharp depression on the size of the secretory peaks. Note that Al^3+^ induced no secretion in the absence of stimulatory pulse. No modification of the basal secretion was observed in the absent of K^+^ pulses (interpulse period) when Al^3+^ was applied. Panel E shows the normalized time course summarizing catecholamine release blockade, after different Al^3+^ concentrations were administered. Thus, in five experiments of three different cultures, the catecholamine release obtained in the presence of Al^3+^ (30, 100, 300 and 1000 μM) amounted to 31.7 ± 5; 44.8 ± 9; 70.4 ± 7 and 89.1 ± 7%, respectively, compared with the pulse P4, immediately before the introduction of Al^3+^. In summary, the Al^3+^ shown depression of catecholamine secretion in a concentration-dependent manner, being the IC_50_ 89.12 μM, as shown on panel F.

### 3.2. Al^3+^ decreases the [Ca^2+^]_c_ level

Previous experiments have shown the drastic depression of catecholamine secretion exerted by the application of Al^3+^. This fact leads us to think that the calcium necessary to activate the secretory machinery may be affected. This has led to design of the experiments represented in Fig. 2. The measurements of the changes in [Ca^2+^]_c_ in populations were carried out by using the fluorescent probe Fluo-4 AM. After 10 s stimulus with K^+^ (35 mM) was applied, which produced a rise of the fluorescence that is maintained during the rest of the experiment that goes on for a minute. Panel A shows original recordings obtained in the absence (control) and in the presence of Al^3+^ (30, 100 and 300 μM). The application of Al^3+^ without cellular stimulation shows no enhancement of fluorescence signal (data not shown). Panel B shown the mean values of the [Ca^2+^]_c_ depression exerted by increasing concentrations of Al^3+^. Al^3+^ 30 μM decreased the fluorescence 13.9 ± 2%; Al^3+^ 100 μM 15.1 ± 2% and the application of 300 μM Al^3+^ exerted the maximal depression of 25.5 ± 2%. This effect in the calcium-dependent fluorescence signal becomes statistically significant with all concentrations of Al^3+^ tested.

### 3.3. Time- and concentration-dependent blockade of I_Ca_ by Al^3+^

Attending that Al^3+^ exerted a depression of neurotransmitter release by a decrease of cytosolic calcium level, we wonder about the origin of this calcium; and one possibility arise from the calcium influx throughout voltage dependent calcium channels. So, in the experiments of Fig. 3 each individual voltage-clamped chromaffin cell tested was stimulated intermittently with 10 ms depolarizing pulses to 0 mV, applied at 10 s intervals from a holding potential of -80 mV. The inward calcium currents (*I*_*Ca*_) were elicited with 10 mM extracellular Ca^2+^ as the charge carrier. In 14 cells tested, the averaged current amounted to 616 ± 59 pA. The current was tested during approximately 10 min, and it suffered no appreciable decline. If a tendency to decline was observed, the cell was discarded. Once the initial current was stabilized, each cell was superfused with different concentrations of Al^3+^, until the effect stabilized. Panel A exhibits an example of the time courses of *I*_*Ca*_ inhibition by three accumulative concentrations of Al^3+^ (100, 300 and 1000 μM). The inhibition exerted by Al^3+^ develops slowly in a concentration dependent manner. In addition, the blockade is irreversible, and the current did not recover after wash-out. Inset shows representative calcium current traces corresponding to control conditions and after perfusion with Al^3+^ (1000 μM). The concentration response curve of the blocking effect of Al^3+^ on peak *I*_*Ca*_ measured at the end of the superfusion period with each Al^3+^ concentration is shown on panel B. Al^3+^ produces a decrease of *I*_*Ca*_, that becomes significant at 300 μM (12.6 ± 2 %) and 1000 μM (61.7 ± 11 %). Al^3+^ shown depression of voltage dependent calcium currents in a concentration-dependent manner, showing a IC_50_ value of 560 μM.

A set of voltage-clamped bovine chromaffin cells was stimulated with increasing strength depolarizing pulses (10 ms), given every 10 s, from –60 mV to +50 mV in 10 mV intervals, from a holding potential of -80 mV, before and 2 min after perfusion with Al^3+^ (1000 μM), using 10 mM Ca^2+^ as charge carrier. *I*_*Ca*_ current-voltage (I-V) relationship shows that in control conditions, I_Ca_ presented a threshold of activation at -40 mV, peaked at around 10 mV (-686 ±59 pA) and had a reverse potential near +50 mV. After 2 min of Al^3+^ (1000 μM) application, peak *I*_*Ca*_ was inhibited to -284 ± 43 pA, and the I-V relationship was shifted towards more positive voltages in which the current activation changed from -40 to -30 mV, but no modifications in the voltage responsible for the maximum peak and current activation-deactivation-kinetics were observed. Al^3+^ inhibited *I*_*Ca*_ more at negative potentials than the positive potentials (60% *vs* 30% approximately) (Panel C).

### 3.4. Effects of Al^3+^on voltage-dependent Ca^2+^ channel subtypes

The different calcium channel subtypes have different physiological roles (García et al., 2006); therefore, it is crucial to study the effect Al^3+^ has on each individual subtype. This can be achieved by using selective calcium channel blockers. In bovine chromaffin cell, L type calcium channels represents around 15% of the total calcium current, whereas N and P/Q represent 30 and 50% respectively; the remaining current corresponded to R-type calcium channels (Hernandez et al., 2011). In next experiments, the chromaffin cells were voltage clamped from a holding potential of -80 mV and stimulated with depolarizing pulses to 0 mV for 10 ms at 30 s intervals. After control current was stabilized, using 10 mM Ca^2+^ as a charge carrier, the cells were firstly superfused during 360 s with ω-conotoxin-GVIA (1 μM, selective irreversible blocker of N-type calcium channels) and ω-agatoxin-IVA (1 μM, selective irreversible blocker of P/Q-type calcium channels). As shown Fig. 4 (panel A), these blockers exerted a reduction of *I*_*Ca*_ of 81.47 %. Then, addition of Al^3+^ (300 μM) during 3 min, a further 9.17 % reduction of the current was observed. The remaining current was blockade with the application of Cd^2+^ (200 μM), and selective and unspecific blocker of calcium channels. No recovery of the current after wash-out was detected. In panel B, after control current was stabilized, the bovine chromaffin cells were superfused during 120 s with nifedipine (3 μM, selective reversible blocker of L-type calcium channels). Nifedipine produced a limited blockade of 12.43% of the peak *I*_*Ca*_, while the addition of Al^3+^ (300 μM) during 60 s, reduced the current an additional 29.65 %. Cd^2+^ (200 μM) was superfused leading to a full blockade of the remaining calcium current.

The degree of I_Ca_ inhibition on the different calcium channels subtypes induced by Al^3+^ was tested at different holding potentials. Panels C-D show the rate and extent of *I*_*Ca*_ blockade by Al^3+^ in different sets of experiments whose holding potentials were initially fixed at -80 mV and after stabilization, changed to -50 mV. Note a loss of 75% of the calcium currents by the inactivation of these channels exerted by the more depolarizing voltage -50 mV. After stabilization of the current (10 mM Ca^2+^), the application of 50 ms test pulses to 0 mV (20 s intervals) generated an *I*_*Ca*_ of 700 pA and decrease to around 125 pA, at -80 and -50 mV holding potential, respectively. In panel C, the administration of nifedipine (3 μM) lead to a full blockade of the recorded current al-50 mV holding potential and no additional depression was exerted by Al^3+^ (300 μM) or Cd^2+^ (200 μM) application. The calcium current was not recovery after wash-out. Panel D shows a similar experiment in which, at -50 mV holding potential, the initial addition of Al^3+^ (300 μM) exerted a full blockade of the remaining current and no effect was observed after perfusion with nifedipine (3 μM) or Cd^2+^ (200 μM). No recovery of the current was observed after wash-out. Note that in this experimental condition, the depolarizing holding potential lead to drastic inactivation of non-L calcium channels subtype, and the remaining L-type current was fully blockade by Al^3+^ in an irreversible manner.

### 3.5. Al^3+^exerts a concentration dependent blockade of I_Na_ currents

Sodium currents are the responsible for the generation and propagation of action potentials firing. In the experiments of Fig. 5, each individual voltage-clamped cell tested was stimulated with 10 ms depolarizing pulses at -10 mV, applied at 30 s intervals from a holding potential of -80 mV. The average initial sodium current (*I*_*Na*_) amounted to 1060 ± 234 pA (n=10). This current suffered no appreciable decline during the experiment; as before, if a tendency to decline was observed, the cell was discarded. Once the initial current stabilized, each cell was perfused with cumulative concentration of Al^3+^ for a 1-min period. Panel A shows the time courses of *I*_*Na*_ with Al^3+^ concentrations of 100, 300 and 1000 μM) applied during the time indicated by the top horizontal bar. These concentrations exerted a fast depression of the *I*_*Na*_ current (10 μM, 8.2 ± 1%; 100 μM, 15.6 ± 3%; 300 μM, 24.3 ± 4% and 1000 μM, 61.1 ± 7%; n=4-8). Furthermore, this blockade is reversible, since, after wash-out, the current was recovered up to a 44.8 %. Inset shows original current traces obtained before (a) and after (b) superfusion with Al^3+^ (1000 μM). Al^3+^ shown depression of voltage dependent sodium currents in a concentration-dependent manner, showing a IC_50_ value of 419 μM (see panel B).

On the other hand, on panel C, a set of voltage-clamped bovine chromaffin cells were stimulated with depolarizing pulses (10 ms) of increasing strength, giving every 10 s, from -60 to +40 mV in 10 mV intervals, from a holding potential of -80 mV, before and after 2 min perfusion of Al^3+^ (1000 μM). *I*_*Na*_ was significantly inhibited, from -791 ± 106 pA in control conditions, to --262 ± 44 pA after 2 min of Al^3+^ perfusion. In this condition, the I-V relationship shifted slightly towards more positive voltages. I_Na_ presented a threshold of activation at -60 mV in both conditions; but peaked at -10 mV and 0 mV in control conditions and after Al^3+^ perfusion, respectively; and also, showed a reverse potential at +40 mV in control versus +30 mV after Al^3+^ administration.

### 3.6 Different effects of Al^3+^ on potassium current subtypes

In bovine chormaffin cells, as in most cell types, K^+^ is the ion responsible for phenomenon of repolarization and termination of action potential firing (Solano et al., 1995). The K^+^ current is associated to two types of channels, those being, voltage dependent K^+^ channels (*I*_*Kv*_) and Ca^2+^/voltage-dependent K^+^ channels (*I*_*KvCa*_) (Platt et al., 1995). In experiments outlined on Fig. 6 the effects of Al^3+^ exclusively on voltage-dependent potassium currents (*I*_*Kv*_) were tested. *I*_*Kv*_ was elicited by 100 ms depolarizing pulses of +120 mV from a holding potential of -80 mV applied a 30 s intervals. 200 μM of Cd^2+^ was added to the extracellular solution to prevent the activation of Ca^2+^/voltage-dependent potassium channels. This voltage-dependent K^+^ current amounted to 2.2 ± 0.5 nA (n=8). Once the initial current stabilized, each chromaffin cell was superfused with cumulative Al^3+^ concentrations (100, 300 and 1000 μM). Low Al^3+^ concentrations (100, 300 μM) did not exert important changes on voltage-dependent potassium currents (2.5 ± 1 and 13.2 ± 2 %, respectively). Nonetheless, at higher concentrations (1000 μM), this metal triggered a mild blockade of the *I*_*Kv*_ (26.2 ± 8.4 %). Panel A shows original traces of *I*_*Kv*_ in control condition and after cell perfusion with 300 μM Al^3+^. Al^3+^ inhibited *I*_*Kv*_ in a concentration-dependent manner, showing a IC_50_ value of 340 μM, as shown on panel B. This blockade was partially reversible, since it slightly recovered after wash-out (data not shown).

The K^+^ current in the chromaffin cell is mainly associated with Ca^2+^/voltage-dependent K^+^ channels (Marty and Neher, 1985). It has been suggested that Ca^2+^ channels and K^+^ channels are in close proximity (Prakriya and Lingle, 1999, 2000). To record *I*_*KvCa*_, cells were initially depolarized with a prepulse to 0 mV (30 ms) to open Ca^2+^ channels and trigger calcium influx; then, a depolarization step to +120 mV (700 ms) was applied (Vm −80 mV) to activated the K^+^ channels (see protocol in panel 7A). The brief depolarizing prepulse elicited a subplasmalemmal calcium rises in the vicinity of the Ca^2+^ channels rather than altering the total cytosolic calcium concentration. At +120 mV (near the equilibrium potential for Ca^2+^), Ca^2+^ influx ceases, but subplasmalemal level of calcium remain, and *I*_*KvCa*_ current activates. Note, that in the absence of the 10-ms prepulse, no Ca^2+^ current is activated, and all the outward current recorded arises from Ca^2+^-independent voltage activated K^+^ channels (*I*_*Kv*_). The amplitude of the outward current activated at +120 mV (without the prepulse) is equal that recorded at +120 mV after the prepulse but preventing the Ca^2+^ load as account in the presence of the calcium channel blocker Cd^2+^ (data not shown). The *I*_*KvCa*_ decay after the closure of Ca^2+^ channels in a rapid and complete manner due to the loss of intracellular Ca^2+^ concentration by intracellular calcium buffers and Ca^2+^ extrusion mechanisms (Roberts, 1993; Naraghi and Neher, 1997). Using this prepulse protocol, the effects of Al^3+^ in the activation of *I*_*KvCa*_ were tested. Fig. 7 shows the inhibition by three concentrations of Al^3+^ (100, 300, and 1000 μM). Al^3+^ was found to reduce *I*_*KvCa*_ by 4.5±2% at 100 μM, 17.9±3% at 300 μM and 45.3±4 % at 1000 μM from an initial *I*_*KvCa*_ of 4.1±0.5 nA. At the end of the wash-out period, both peak and late current reached a similar amplitude of the *I*_*KvCa*_ recording during the Al^3+^ perfusion.

This suggests that most of *I*_*KvCa*_ blockade exerted by this metal could account for blockade of Ca^2+^ entry through voltage-dependent calcium channels. Panel A shows *I*_*KvCa*_ original traces corresponding to control conditions (a) and after perfusion (b) with Al^3+^ (1000 μM). Inhibition of *I*_*KvCa*_ was measured at the end of the 2-min superfusion period with each Al^3+^ concentration and the IC_50_ value was 447 μM (see concentration response curve shown in panel B).

The activation of Ca^2+^/voltage dependent K^+^ channels can be identified by a characteristic hump in the intensity-voltage (I-V) curve (see panel 7C). The effects of Al^3+^ on *I*_*KvCa*_ were examined on I-V curve recordings. When Ca^2+^ influx is prevented by 100 μM Cd^2+^ application, a large fraction of K^+^ outward current is inhibited and the hump in the I-V relationship disappears (data not shown). Comparison of the I-V relationship in the presence and absence of Ca^2+^ influx indicates that in this cells, most K^+^ outward current is Ca^2+^/voltage dependent, in especial, at the voltage range values associated with the action potential (-30 to +50 mV). Panel C shows the effects of Al^3+^ (300 μM) on outward calcium and voltage dependent K^+^ currents. Panel shows the peak of outward currents activated by depolarization steps (700 ms duration) repeated at 10 s intervals from a -80 mV holding potential with +10 mV increments, in 2.5 mM external calcium. *I*_*KvCa*_ shows a threshold of activation at around -30 mV and the peak current was detected at +30 mV with a value of 2223 pA. As indicated, this relationship exhibited a pronounced hump. This hump appeared to mirror the I-V relationship of the Ca^2+^ current in these cells (see Fig. 3). After 2 min application of Al^3+^ (300 μM), a substantial depression in the peak *I*_*KvCa*_ was observed. The residual current-voltage relationship displayed a linear behavior. Note the drastic reduction of the *I*_*KvCa*_ amplitude evoked by Al^3+^ at +30 mV, where the contribution of *I*_*KvCa*_ to the total outward K^+^ efflux was higher than at more depolarizing potentials.

### 3.7. Al^3+^induces a depression of evoked action potentials

Using perforated-patch recording and 2.5 mM external Ca^2+^, evoked action potentials (APs) were elicited in chromaffin cells by current injection through the recording electrode (Fig. 8). The membrane potential was hyperpolarized to -80 mV to prevent the inactivation of sodium conductance. The current injection needed to trigger a stable train of APs was 10 pA above the resting potential, and the duration of the stimulus was 1000 ms. Only the AP elicited by the depolarizing pulse was used for analysis because bovine chromaffin cells do not fire spontaneously. With higher current injections, the chromaffin cells did not fire at higher frequencies (data not shown). If the current pulse was made longer, the amplitude of the spike decreased with time, and the regular firing was interrupted near the end of the pulse by a progressive decline of spike amplitude, probably due to inactivation of sodium currents caused by sustained depolarization (data not shown). In control conditions, the stimulation of the cell triggered the initiation of approximately 1-2 action potentials every 100 ms (13.7 ± 1.6 APs in 1000 ms, n=13) (see Fig. 8). We observed no detriment on the firing frequency through either the stimulation pulse nor the following pulses. The mean peak amplitude of evoked APs in control conditions was 31.6 ± 3.6 mV with a rise time of 16.2 ± 1 ms. The half-width of the APs was 16.8 ± 1.2 ms and the area 530 ± 10 mV/ms. After APs repolarization (decay time of 7.6 ± 0.7 ms), the afterhyperpolarization (AHP) amplitude was -13.6 ± 1.3 mV and the area -174 ± 27. After 2 min of Al^3+^ perfusion (100 and 300 μM, top panel and bottom panel, respectively) the membrane potential (Vm) became more depolarized (control -65 ± 7 mV; 100 μM, -54.2 ± 4 mV; and 300 μM, -45 ± 3 mV). We observed an increase of resting membrane potential after Al^3+^ administration, that at 100 μM barely had any effect without affecting the action potential firing nor the biophysical properties of APs; but that became significant at 300 μM, where the spontaneous APs were inhibited as the membrane stayed depolarized, showing finally a prominent plateaus phase that correlated to a drastic depression on the action potential firing. The evoked APs became progressively lower and broader with a concomitant decrease in afterhyperpolarization (AHP) amplitude and area (Table 1). These effects could be related to the progressive decrease in AHP duration during the pulse due to the smaller Ca^2+^ influx and the sustained depolarization that lead to Na^+^ current inactivation and the increasing difficulty to generate APs. Note in panel 8B no recovery in amplitude and frequency of APs occurred after Al^3+^ wash-out.

**Table 1.** Kinetic parameters of evoked individual action potential events in different experimental conditions. Al^3+^ administration (100 μM top panel, and 300 μM bottom panel). Data are means ± SEM of 397-523 APs of 6-8 different rats. Statistical analysis was performed by Mann Whitney test. ^*^p < 0.05, ^**^p < 0.01, ^***^p < 0.001 respect to control.

|  | Amplitude<br>(mV) | <sup>10-90</sup>Rise<br>(ms) | <sup>10-90</sup>Slope<br>(mV/ms) | <sup>90-10</sup>Decay<br>(ms) | <sup>90-10</sup>Slope<br>(mV/ms) | Area<br>(mV.ms) | Halfwidth<br>(ms) | AHP<br>(mV) | AHP area<br>(mV.ms) |
| --- | --- | --- | --- | --- | --- | --- | --- | --- | --- |
| Control | 31.6 ± 3.6 | 16.2 ± 1.0 | 1.5 ± 0.1 | 7.6 ± 0.7 | - 3.5 ± 0.9 | 530 ± 66.2 | 16.8 ± 1.2 | - 13.6 ± 1.3 | - 174 ± 27 |
| Al<sup>3+</sup> | 22.7 ± 1.7* | 15.9 ± 0.8 | 1.1 ± 0.1** | 11.6 ± 1.3* | - 1.2 ± 0.2** | 470 ± 62.3 | 19.4 ± 1.5* | - 6.7 ± 0.8*** | - 85.4 ± 16*** |
| Washout | 22.4 ± 2.5* | 14.1 ± 1.7 | 1.5 ± 0.3 | 11.1 ± 1.6* | - 1.6 ± 0.5* | 439 ± 59.9 | 17.8 ± 2.1 | - 7.1 ± 1.0*** | - 86.5 ± 19*** |

## 4. Discussion

In this study, we analysed the effects of acute aluminium (Al^3+^) administration on the excitability of bovine chromaffin cells by analysis of neurotransmitter release, intracellular calcium levels, and calcium, sodium and potassium currents, as well as, on the generation and propagation of evoked action potentials. Thus, Al^3+^ produced (1) a 50% reduction of catecholamine release, (2) a 20% decrease on intracellular Ca^2+^ levels, (3) a gradual and irreversible blockade of voltage-dependent Ca^2+^ currents with a IC_50_ of 560 μM, (4) a gradual and reversible blockade of voltage-dependent Na^+^ currents with a IC_50_ of 419 μM, (5) a few depression on voltage-gated K^+^ currents, (6) a blockade of Ca^2+^/voltage-dependent K^+^ currents with a IC_50_ of 447 μM and, (7) a total suppression of the generation and propagation of action potentials.

Several metals are known to act at synaptic level and have a role in neuronal excitation (Tamano and Takeda, 2011), thus, Pb^2+^ occupies calcium-binding sites on numerous calcium-dependent proteins (Marchetti, 2014) and has a direct effect on both voltage- and glutamate-activated calcium channels (Büsselberg et al., 1994a,b); Cd^2+^ mimics calcium, blocking calcium channels and receptor operated calcium channels (Choong et al., 2014). Ni^2+^ modifies the expression of some genes that codify for ionotropic glutamate receptors, and blocks also calcium currents (Slotkin and Seidler, 2009; Kang et al., 2006, 2007); Mn^3+^ inhibits acetylcholinesterase inducing a characteristic parkinsonian-like syndrome (Santos et al., 2012); Zn^2+^ modulates several ionic conductances (Baraibar et al., 2020, Neumaier et al., 2020; Trombley et al., 1998) and modifies long-term potentiation (LTP) and long-term depression (LTD) (Nakashima and Dyck, 2009; Song et al., 2014); Cu^2+^ modifies NMDA receptor (Schlief et al., 2006) inhibiting both glutamate-mediated neurotransmission (Mathie et al., 2006; Schlief et al., 2005) and LTP (Goldschmith et al., 2005). These actions induce aberrant synaptic neurotransmission that leads to neurological diseases (Tamano and Takeda, 2011).

Compounds containing Al^3+^ are employed in a variety of human-related products and procedures (Scancar and Milacic, 2006), however, the main source of Al^3+^ intake is food due to both its original levels and interactions with the materials employed in its storage and preparation (Krewski et al., 2009, Igbokwe et al., 2020). Although Al^3+^ absorption is minimal, it varies significantly depending on the diet. On the whole, part of the population might be at risk of exceeding the permitted tolerable weekly intake (Fekete et al., 2012) being at risk of developing pathologies related to Al^3+^(Krewski et al., 2009). In fact, it exists a clear association between high levels of Al^3+^ in the CNS and neurotoxicity (Krewski et al., 2009; Kawahara and Kato-Negishi, 2011). Al^3+^ is absorbed (Jaishankar et al., 2014), it reaches systemic blood circulation where is transported (Nagaoka and Maitani, 2005), crosses the blood brain barrier accumulating in brain (Deloncleet al., 1990). Up to day, the main mechanism of Al^3+^ neurotoxicity reported is the reactive oxygen species production, giving rise to oxidative stress and mitochondrial dysfunction (Kumar and Gill, 2014). Al^3+^ produces a shift in brain homeostasis towards increased proinflammatory cytokine production (Johnson and Sharma, 2003), while also acting as a pro-oxidant (Exley, 2004). In addition, Al^3+^ binds to important antioxidant enzymes as superoxide dismutase altering its activity (Di et al., 2006). The Al^3+^-superoxide complex aggravates oxidative damage enhancing the redox activity of iron and elevating the concentration of important bio-oxidants like the OH radical (Ruipérez et al., 2012). Al^3+^ also enhances NADH peroxidation (Exley, 2004) and increases iron-induced peroxidation in liposomes (Kaneko et al., 2014) and erythrocytes in a concentration-dependent manner (Gutteridge et al., 1985).

Al^3+^ seem to be able of altering long-term potentiation (LTP), affecting synaptic plasticity and, thus, memory (Li et al., 2020; Wang et al., 2010). In fact, Al^3+^ interferes with phosphorylating processes affecting cAMP (Johnson et al., 1987), alters Ca^2+^/calmodulin-dependent enzymes (Levine et al., 1990), interferes with G-proteins (Haug et al., 1994; Shafer et al., 1993); all of them, targets involved in LTP process (Harris et al., 1984; Frey and Kandel, 1993; Malenka et al., 1989, Matthies and Reymann, 1993). Moreover, Al^3+^ alter calcium influx (Koenig and Jope, 1987), modifies the calcium binding to calmodulin (Siegel and Haug, 1982), and displaces calcium from phospholipid binding sites (Farnell et al.,1985). These data combined with the decreased activity of AChE induced by Al^3+^ (Yellamma et al., 2010), may be responsible for the abovementioned memory impairments (Martinez et al., 2017). Additionally, Al^3+^ may alter mRNA levels of IL-1β, IL-6, TNF-α, MCHII, CXC3CL1 and BDNF, leading to a neuroinflammatory process that is going to further aggravate memory deficits (Cao et al., 2016).

It has been reported that Al^3+^ binds to calcium channels (Busselbert et al., 1994) modulating the amplitude of excitatory post-synaptic potentials (Platt and Büsselberg, 1994; Platt et al., 1995); while its effects on other ionic currents are less remarkable (Meiri and Shimoni, 1991). Therefore, in this study we have observed a drastic reduction of neurotransmitters release exerted by Al^3+^ in a concentration dependent manner, up to an 86 % at high concentrations, being the IC_50_ 89.12 μM (see Fig. 1). This may be related to the property reported that Al^3+^ may alter hippocampal neuronal activity (Farnell et al., 2013), may reduce the amplitude of action potentials (Platt et al., 1995) and may modulate voltage-gated calcium channel currents (Büsselberg et al., 1994). However, attending other studies, where it has been reported that Al^3+^ lead to an elevation on calcium concentration by calcium released from intra-axonal organelles or calcium-binding proteins; in combination with the impaired uptake of calcium due to the interaction between Al^3+^ and Mg^2+^-dependent ATPase (Platt et al., 1995; Siegel and Haug, 1983a, b), a potentiation of secretions may be expected. However, in our hand, Al^3+^ also reduces intracellular calcium concentrations around a 25% (see Fig. 2) in accordance with a reduction of neurotransmitter release.

**Fig. 1.**
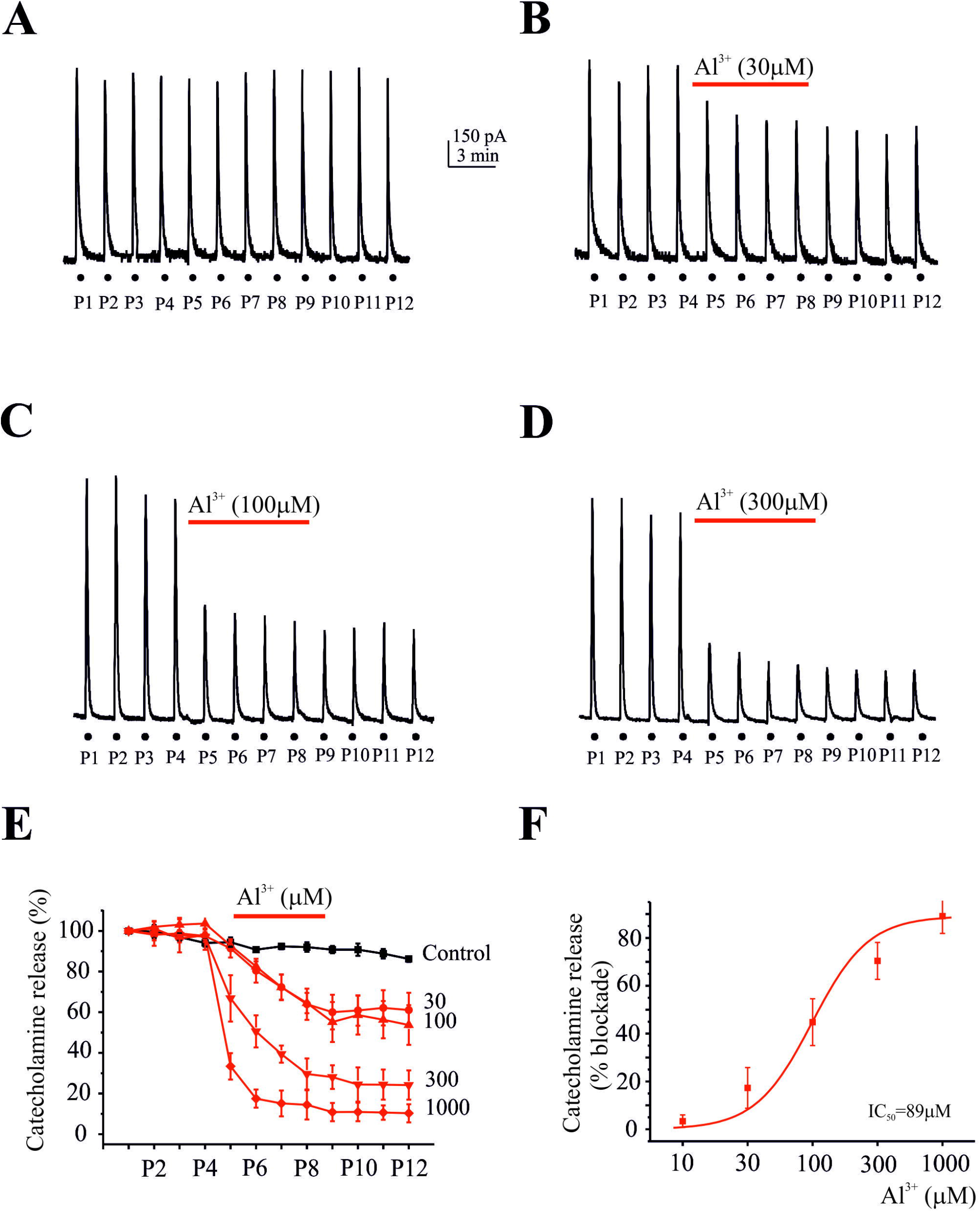
Al^3+^ depress the catecholamine release responses obtained in fast-superfused cells triggered by K^+^ stimulation. Cells were superfused with Krebs-HEPES solution containing 2 mM Ca^2+^ and stimulated at 3 min intervals with 5 s pulses of 35 mM K^+^. Al^3+^ was applied as indicated by the top horizontal bars. **A** Original records obtained in control cells. **B**-**C**-**D** Original traces after administration of Al^3+^ at the concentrations indicated. **E** Averaged time course of the effects exerted by different concentrations of Al^3+^. Data points expressing % of neurotransmitter inhibition (ordinate axis), after different Al^3+^ concentrations were used (red points) in comparison to control (black points). A separate cell population was used for each individual concentration. **F** Sigmoidal Hill function [*y*=start + (end-start) *x*^n^/(*k*^n^ + *x*^n^)] fitted with the averaged data of the % of catecholamine release inhibition (ordinate axis) after administration of each Al^3+^ concentration (abscissa axis). IC_50_ was 89 μM. The data plots were normalized to the mean value of the control period and expressed as the mean ± SEM of 4-5 experiments.

**Fig. 2.**
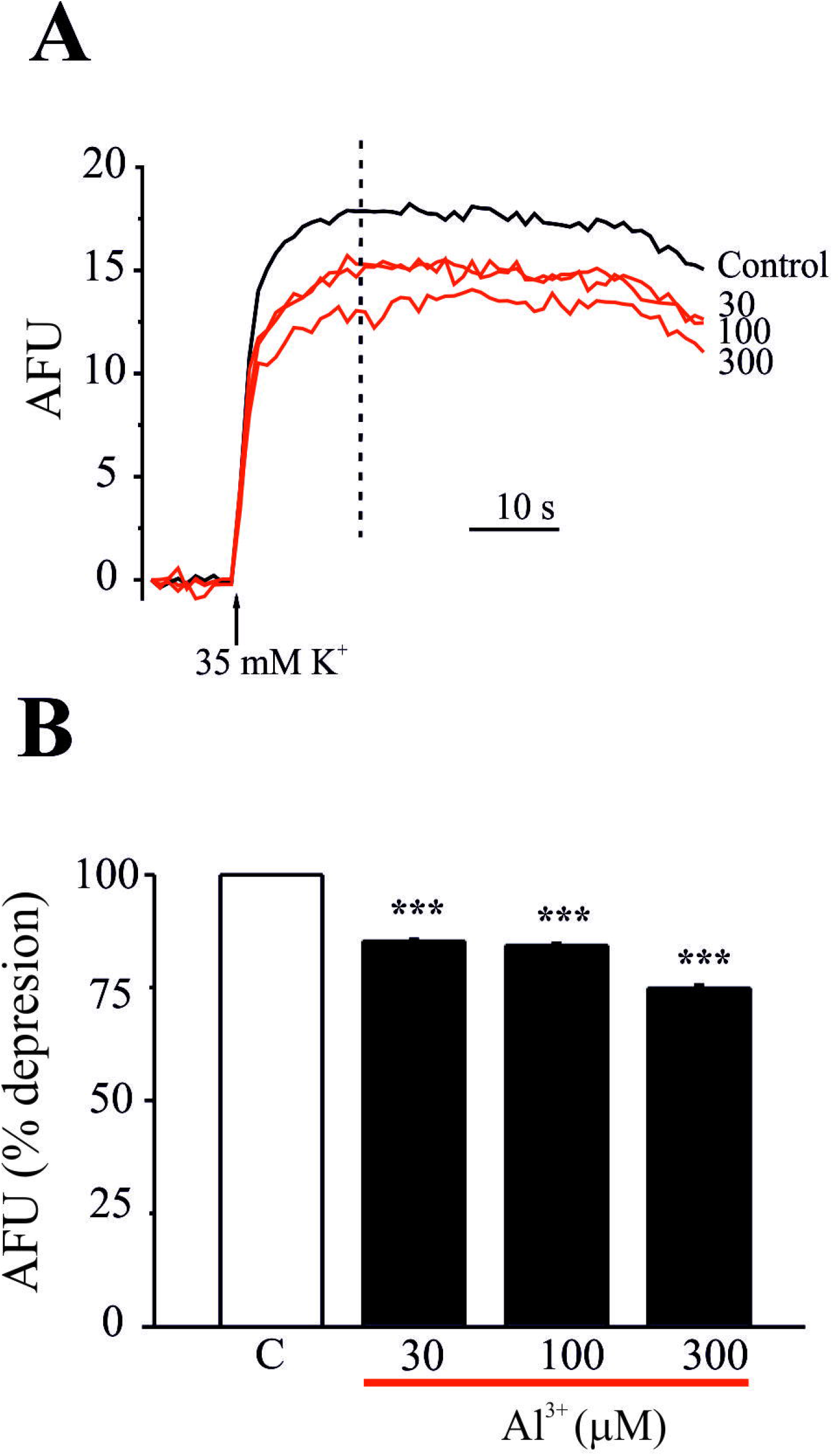
Effects of Al^3+^ on intracellular Ca^2+^ levels. **A** Original traces of the Fluo-4 fluorescence induced by 35 mM K^+^ in the absence (control conditions, black line) or presence of Al^3+^ at the indicated concentrations (red lines). **B** Bar diagram comparing the percentage of the effect in fluorescence in the absence (control) or presence of Al^3+^ at concentrations indicated in abscissa. The values have been normalized with the maximum fluorescence increment produced with K^+^ (35 mM). Data expressed as the mean ± SEM of 8-9 experiments from 3 different batches of cells cultures. \*\*\**p* < 0.001 respect to control. AFU: Arbitrary Fluorescence Units.

Calcium play an essential role in neurotransmitter release (Katz and Miledi, 1968) due to their influx through the voltage-dependent calcium channels (Miledi, 1973), essentially through N- and P-types in neurons (Takahashi and Momiyama, 1993) and L-type in neuroendocrine cells (Marcantoni et al., 2007). Primary cultures of chromaffin cells from the bovine adrenal medulla express L-type (15%), N-type (35%) and P/Q-type (50%) calcium channel subtypes (García et al., 2006). Additionally, also a R-type channels may be observer under specific conditions (Hernandez et al., 2011). Some studies have reported an opposite effect of Al^3+^ on voltage dependent calcium channels, at low concentrations Al^3+^ triggers a use-dependent and irreversible blockade of Ca^2+^ currents (Meiri and Shimoni, 1991; Platt and Büsselberg, 1994) while high levels of Al^3+^ increase these conductance (Chen et al., 2005). It seems that Al^3+^ may binding to at least three different specific binding sites in calcium channels, generating a selective effect on calcium channels, with less action on voltage-activated sodium and potassium channel currents (Busselberg et al., 1994). Furthermore, we showed that Al^3+^ blocks Ca^2+^ currents in a time- and concentration-dependent manner and that this blockade is irreversible (see Fig. 3), stablishing an IC_50_ of 560 μM. Nevertheless, we were unable to observe that higher levels of Al^3+^ increase Ca^2+^currents. The highest concentrations used (1000 μM), produced a 62% blockade of Ca^2+^ currents, implying that all Ca^2+^ channel subtypes present on bovine chromaffin cells are affected by this metal. Nonetheless, Ca^2+^ currents were more inhibited at negative potentials. Al^3+^ slightly shifted the I-V relationship towards more positive voltages, with no modifications on its kinetic parameters, suggesting that the blockade occurred regardless of the opening or closing of channels and that there exists a partial selectivity of Al^3+^ towards one Ca^2+^ channel subtype. L-type are activated at more negative potentials than the N- and P/Q-calcium channel subtypes. Hence, it appears reasonable to believe that Al^3+^ effect is voltage dependent. On the bovine chromaffin cell, there exists a bigger proportion of N- and P/Q-calcium channel subtypes (81%) than L-subtype (15%) (see Fig. 4), thus, Al^3+^ was noted to have a slight difference in its affinity, showing a higher blockade on ω-toxin-resistant currents (L types) (50%) than nifedipine-resistant currents (N- and P/Q types) (35%). A rate of both, the blockade exerted by Al^3+^ and also the remaining current, may be associated to the R-type Ca^2+^ current that may contribute, as previously reported in rodent chromaffin cells, to around 15 % of the total Ca^2+^ current (Hernandez et al., 2011; Hernández-Guijo et al., 1998; Marcantoni et al., 2010). This action on calcium channels, in combination with the selectivity of Al^3+^ for neuronal tissue, could have significant neurotoxic implications because calcium influx is essential for proper neuronal function, L channels are involved in gene expression whereas N and P/Q channels are responsible for neurotransmission (Baldelli et al., 2000). Additionally, during action potential firing, calcium currents are involved in both, the early slowly activating phase (prespike) carried by L-type channel that contributes to the pacemaker potential and the rapid action potential upstroke; and in the late short-lasting component (postspike) carried by non-L type channels that sustains the AP repolarization (Marcantoni et al., 2010). In summary, the interaction of Al^3+^ with voltage-dependent calcium currents disrupts neuronal calcium homeostasis (Koenig and Jope, 1987; Farnell et al., 1985), which in turn alters neuronal transmission.

**Fig. 3.**
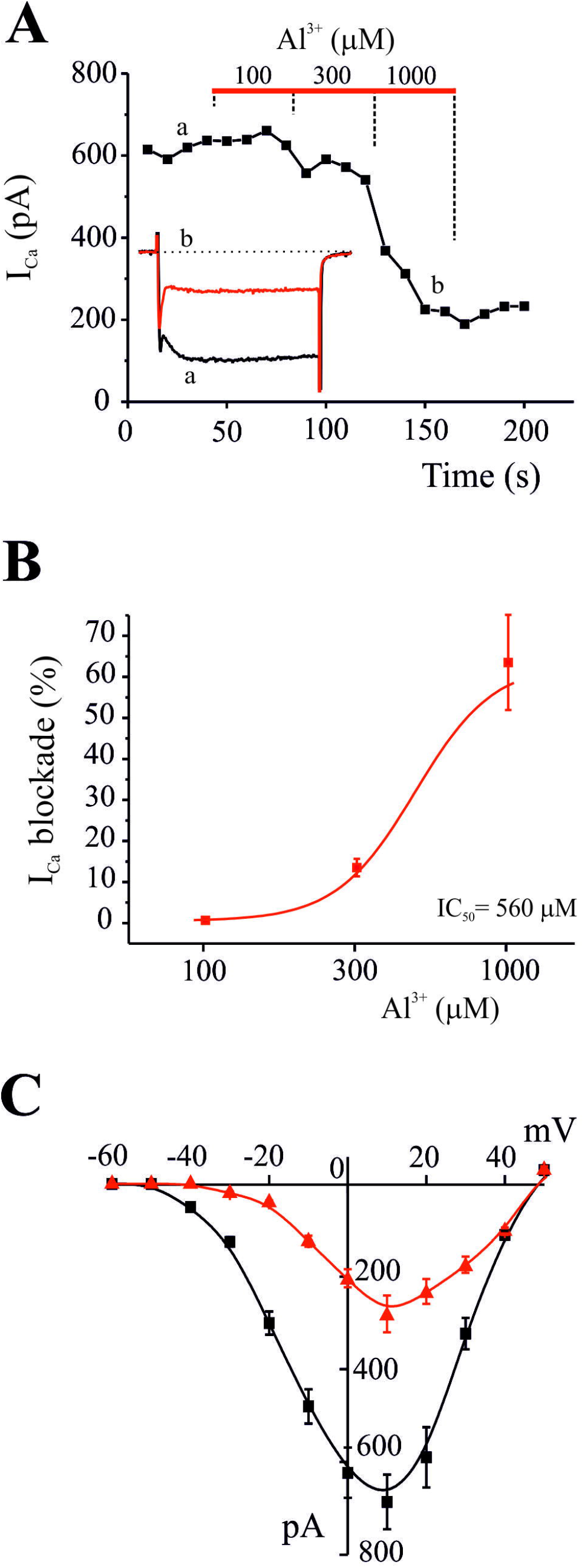
Inhibition of the whole-cell inward Ca^2+^ current exerted by Al^3+^. **A** Representative time course of peak calcium current in control conditions and during application of accumulative concentrations of Al^3+^ during the period indicated by the top horizontal bar. Inset shows original traces obtained from control conditions (a) and after Al^3+^ (1000 μM) administration (b). **B** Averaged results of the % of current inhibited (ordinate axis) after superfusion with each Al^3+^ concentration (abscissa axis). IC_50_ was 560 μM. The data plots were normalized to the mean value of the control. **C** Voltage/Ca^2+^-current relationship obtained before and after perfusion with 1000 μM Al^3+^. Depolarizing pulses were applied at the indicated voltages (abscissa axis); the averaged currents generated are represented on the ordinate axis in control conditions (black line, square symbols) and after perfusion with Al^3+^ (red line, triangle symbol). The data plots were expressed as the mean ± SEM of 7-8 experiments.

**Fig. 4.**
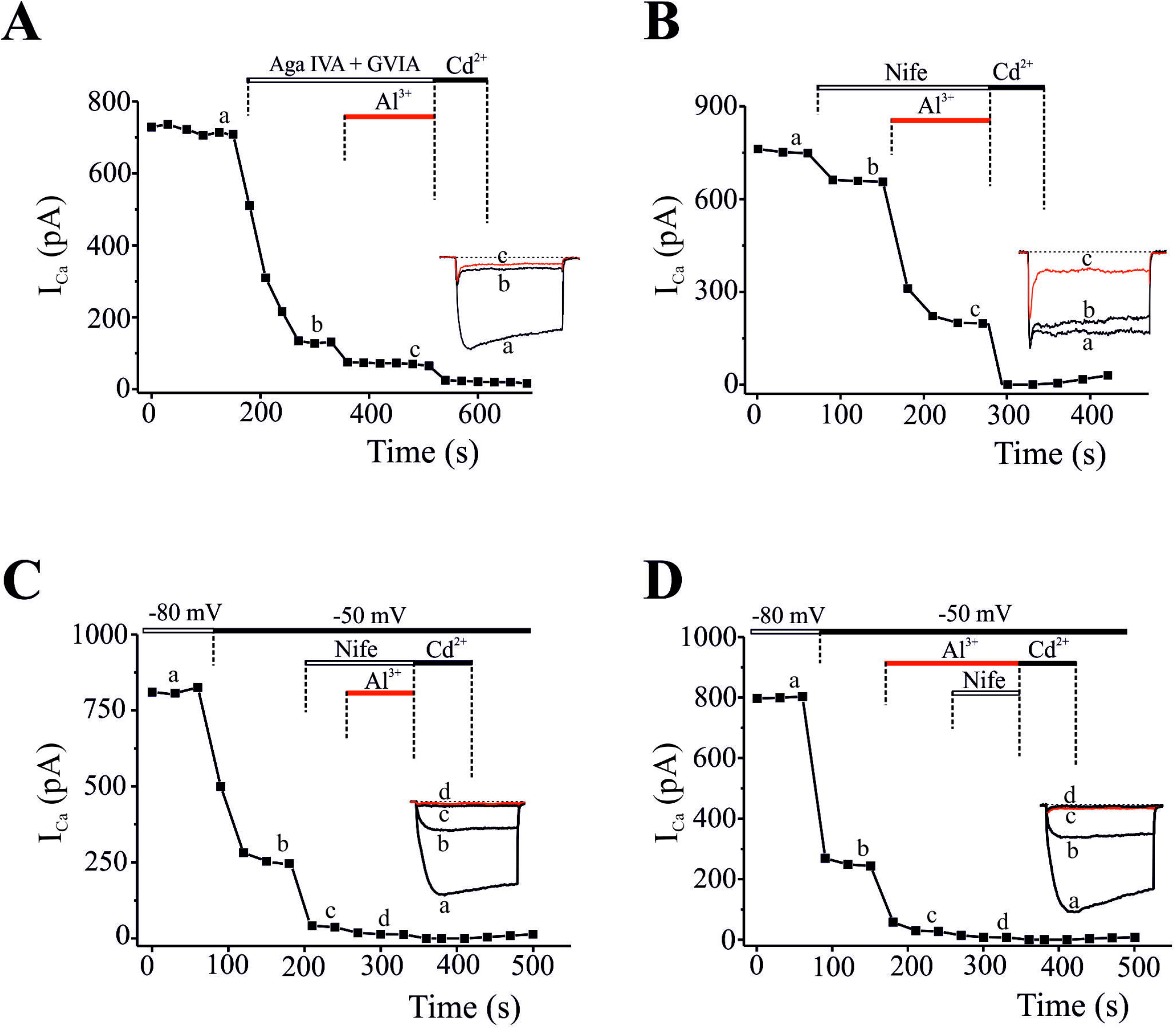
Effect of Al^3+^ on different Ca^2+^ channels subtypes. **A** Time course of the blockade of *I*_*Ca*_ after sequential addition of the selective blocker of P/Q type calcium channels (ω-agatoxin IVA 1 μM) plus the selective blocker of N-type (ω-conotoxin GVIA 1 μM) and Al^3+^ (300 μM) during the time indicated by the top horizontal bars. **B** Time course of the blockade of *I*_*Ca*_ after sequential addition of the selective blocker of L-type (nifedipine 3 μM) and Al^3+^ (300 μM) (top horizontal bars). Insets show original traces at time indicated by letters. **C**-**D** *I*_*Ca*_ blockade by Al^3+^ at -50 mV holding potential. The cells were clamped at -80 mV holding potential and them changed to -50 mV (note drastic loss of current by voltage inactivation of calcium channels). After stabilization of the current, nifedipine (3 μM) and Al^3+^ (300 μM) were applied during the time indicated in the top horizontal bars. Insets show original traces at time indicated by letters. The calcium channels blocker Cd^2+^ (200 μM) was added at the end of the experiments. The data shown were expressed as the mean ± SEM of 5-6 experiments.

In most excitable cells, the input current that triggers the action potential is produced by voltage-dependent Na^+^ channels (Hodgkin and Huxley, 1952). Although various studies have been incapable of demonstrating significant effects of Al^3+^ on Na^+^ channels (Meiri and Shimoni, 1991; Platt and Busselberg, 1994), in our hands, Al^3+^ produced a reversible and concentration-dependent blockade of voltage-dependent Na^+^ currents, reaching a 61% inhibition, and stablished an IC_50_ at 410 μM (see Fig. 5). Moreover, this inhibition was more evident at more negative potentials, the I-V relationship shifted towards more positive voltages. In the majority of excitable cells, the entrance of Na^+^ to the cell, previous activation of voltage-dependent Na^+^ channels, is necessary for the triggering of action potentials (Hodgkin and Huxley, 1952). Hence, the inhibition of Na^+^ channels exerted by Al^3+^ would limit the initiation and progression of action potentials, affecting synaptic neurotransmission and neuronal plasticity.

**Fig. 5.**
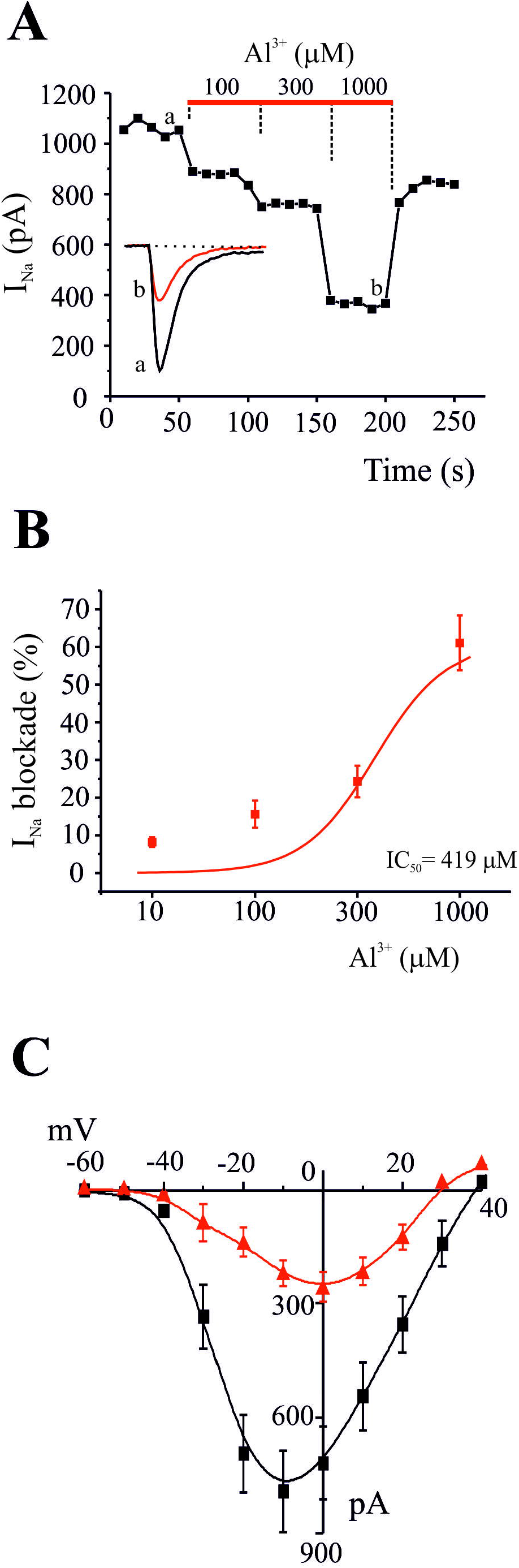
Inhibition of the whole-cell inward Na^+^ currents by Al^3+^. **A** Representative time course of peak sodium current in control conditions and during application of accumulative concentrations of Al^3+^ (see top horizontal bar). Inset shows original traces obtained from control conditions (a) and after Al^3+^ (1000 μM) administration (b). **B** Averaged results of the % of current inhibited after superfusion with each Al^3+^ concentration (abscissa axis). IC_50_ was 419 μM. The data plots were normalized to the mean value of the control. **C** Voltage/current relationship obtained before and after perfusion with 1000 μM Al^3+^. Depolarizing pulses were applied at the indicated voltages (abscissa axis); the sodium currents generated are represented on the ordinate axis in control conditions (black line, square symbols) and after perfusion with 1000 μM Al^3+^ (red line, triangle symbol). The data plots were expressed as the mean ± SEM of 9 experiments from 2 different cultures.

K^+^ channels play a fundamental role in the repolarization of the action potential, setting the resting potential, modifying cellular excitability and temporal regulation of the firing pattern of action potentials (Stocker, 2004; Sun et al., 2009). Among the great diversity of K^+^ channels, in the chromaffin cell, the K^+^ current is associated with both voltage-dependent K^+^ and Ca^2+^/voltage dependent K^+^ channels (BK) (Scott et al., 2011). On our experiments, voltage-dependent K^+^ currents suffered no appreciable blockade after Al^3+^ administration (see Fig. 6), in accordance with what other studies have claimed (Meiri and Shimoni, 1991; Platt and Busselberg, 1994). The loss of K^+^ current that occurs in absence of Ca^2+^ or in the presence of Ca^2+^ channel blockers indicates the marked presence of BK channels in chromaffin cells (Albiñana et al., 2015; Marcantoni et al., 2010). BK channels are activated by voltage but drastically potentiated by low [Ca^2+^]_c_, resulting in hyperpolarization of the membrane potential and alterations in cellular excitability. We observed that Al^3+^ produced a concentration-dependent inhibition of Ca^2+^/voltage-dependent K^+^ currents of around a 45% at the higher concentration used (see Fig. 7). This blockade was reversible, and the IC_50_ was stablished at 447 μM. During the I-V recording, we observed a hump of output current, predominant in holding potentials of -40 to +80 mV, due probably to an increase in cytosolic calcium entry through calcium channels (Marrion and Tavalin, 1998; Wisgirda and Dryer, 1994). Therefore, BK channels show co-localization with various Ca^2+^ channels subtypes. BK channels are activated by calcium influx through L-type calcium channels in mouse chromaffin cells (Marcantoni et al., 2010), or L- and Q-type calcium channels in rat chromaffin cells (Prakriya and Lingle, 1999, 2000). In chromaffin cells, as well as in many neurons, calcium influx during action potentials stimulates K^+^ channels, assisting in repolarization and the generation of long-lasting post-hyperpolarization, which regulates the frequency with which action potentials fire. As a result, Al^3+^ effect on calcium influx has a direct impact on the functionality of these potassium channels and, thus, cell excitability and action potential firing. By other hand, the small conductance (SK) K^+^ channels play an important role in setting the intrinsic firing frequency (Strocker, 2004), while BK channels regulate AP shape; and that both are activated by intracellular Ca^2+^; so, the modulation of calcium influx exerted by Al^3+^ may affect the functionality of these potassium channels drastically impacting the AP firing. In fact, the Carbone lab has described a functional coupling between SK and L-type calcium channels in mouse chromaffin cells (Vandeal et al., 2012). So, this effect of Al^3+^ on calcium channels could avoids both the activation of BK channels responsible for the repolarization phases of the AP and the fast phase of afterhyperpolarization (AHP), and presumably, the activation of SK channels responsible for the post-hyperpolarization phases which determines the arrival of the next AP (Akiyama et al., 2010).

**Fig. 6.**
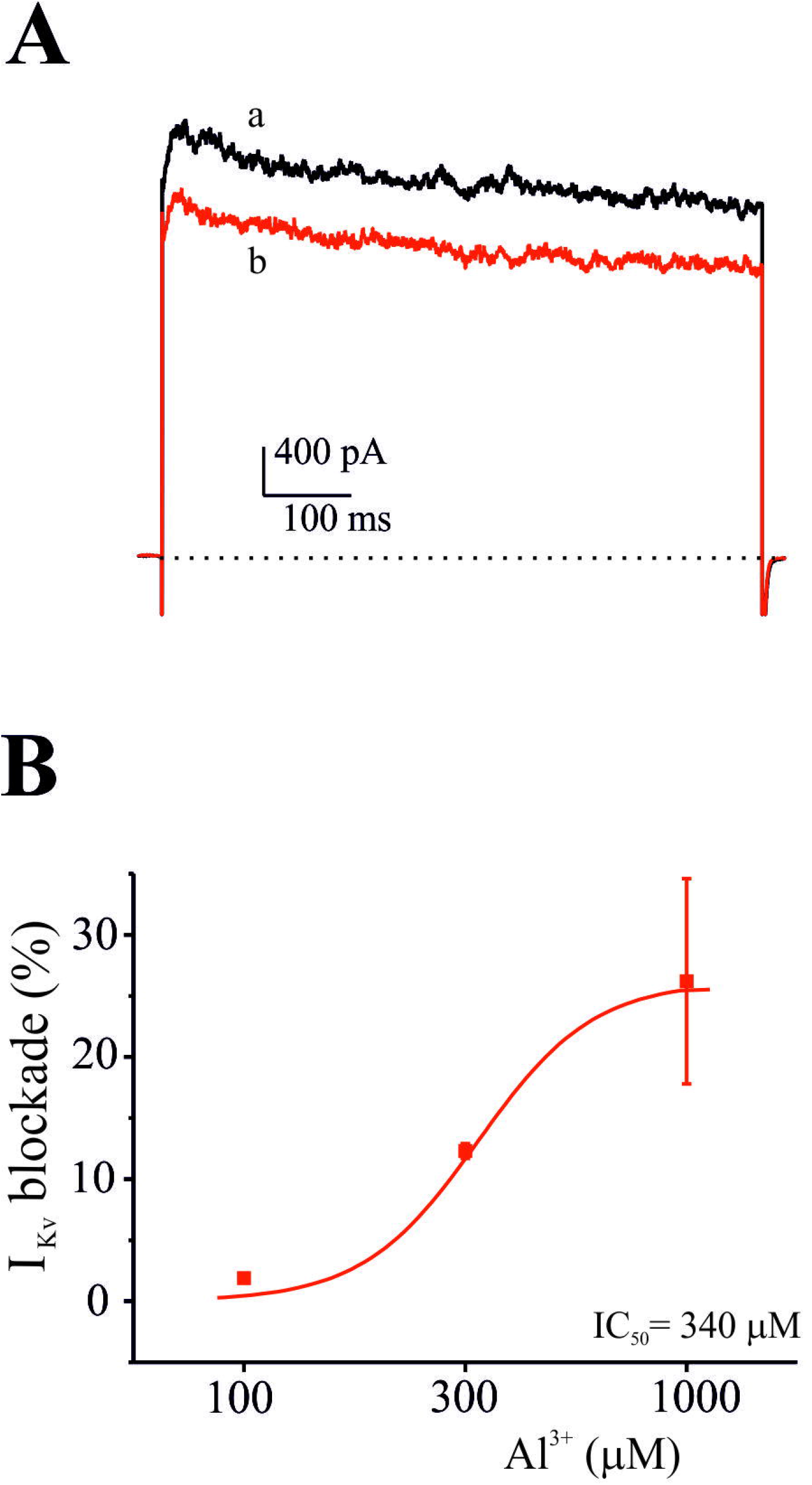
Partial inhibition of voltage-dependent K^+^ currents by Al^3+^. **A** Original traces corresponding to control conditions (a) and 2 min after perfusion with 300 μM Al^3+^(b). **B** Averaged results of the % of potassium current inhibited (ordinate axis) after superfusion with each Al^3+^ concentration (abscissa axis). IC^50^ was 340 μM. The data plots were expressed as the mean ± SEM of 8 experiments.

**Fig. 7.**
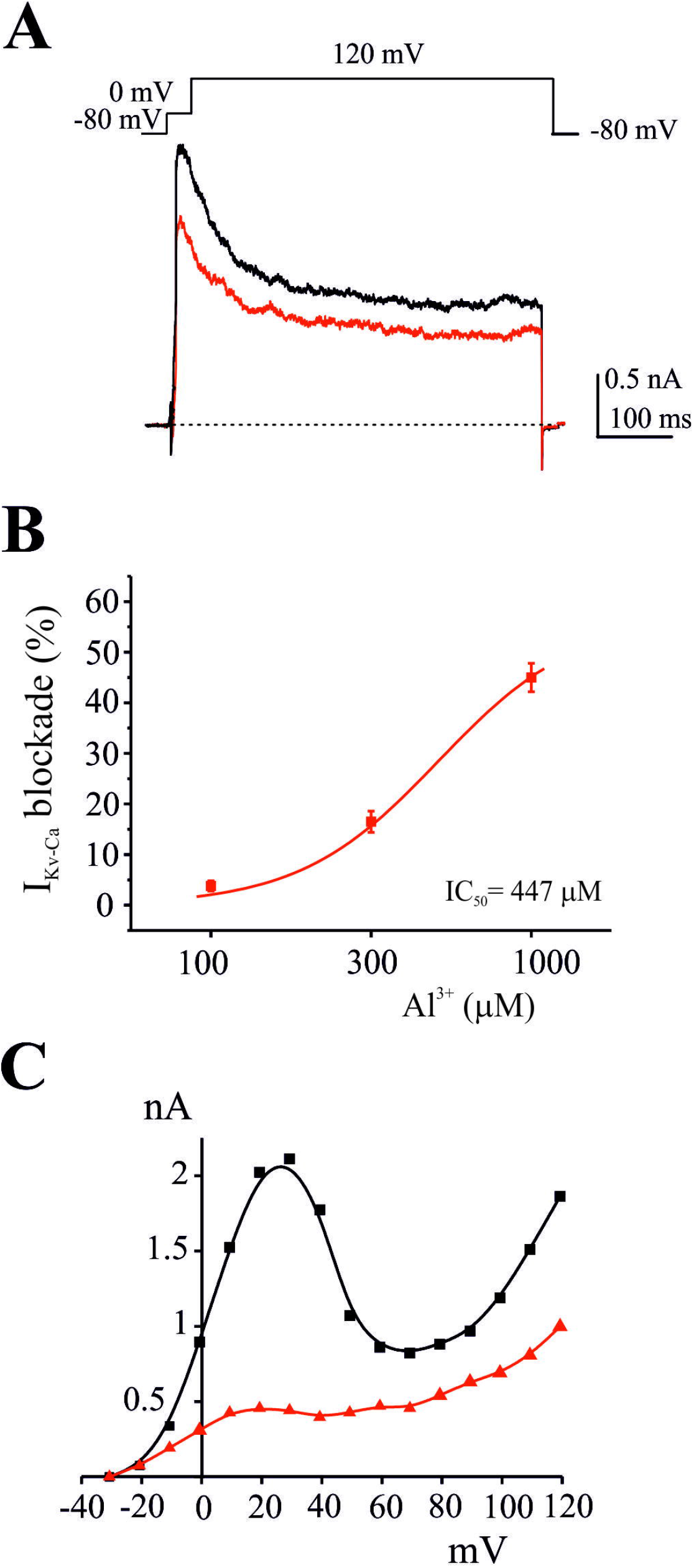
Blockade of Ca^2+^/voltage-dependent K^+^ currents by Al^3+^. **A** Original traces corresponding to control conditions (a) and after perfusion with 300 μM Al^3+^ (b). Note the stimulating protocol used to obtain these outward currents. **B** Averaged results of the % of current inhibited (ordinate axis) after superfusion with each Al^3+^ concentration (abscissa axis). IC_50_ was 447 μM. The data plots were expressed as the mean ± SEM of 8 experiments. **C** Outward K^+^ current versus membrane potential during the test depolarization (I-V plot), before (black line, square symbols) and after perfusion (2 min) with 1000 μM Al^3+^ (red line, triangle symbol). Note than the Ca^2+^/voltage-dependent K^+^ current hump is primarily affected by the exogenous Al^3+^ application.

The inhibition exerted by Al^3+^ on the ionic currents involved in neurotransmitter release (calcium currents), initiation of action potentials (sodium currents) and termination of action potentials (BK currents), further define its neurotoxic nature and may lead to alterations in cellular excitability and synaptic transmission. Whilst it has been reported that this metal could enhance excitability (Platt et al., 1995; Banin and Meiri, 1987; Koening and Jope, 1987), as above indicated, we show that, in addition to a direct depression of neurosecretion, Al^3+^ blocks the main ionic currents responsible for the onset and propagation of action potentials. We show that Al^3+^ depolarized the membrane of the bovine chromaffin cell, decreasing the frequency of APs (see Fig. 8). Thus, the calcium channel blocking results in a reduced functionality of potassium channels involved in repolarization and production of long-lasting post-hyperpolarization, which determines the frequency of action potentials firing, and so, depressing the synaptic transmission. In this way, the mild depolarization of the resting membrane prevents the recovery of sodium channels form inactivation, together with the almost total blockade of the AHP, could explain the decrease in the firing frequency of the APs. Additionally, studies have shown that many molecular mechanisms involved in synaptic plasticity, including protein phosphorylation, gene expression, and neurotransmitter release are regulated by voltage-dependent Ca^2+^ channels (Deisseroth etal., 1998), therefore there exists an intimate relationship between long-term potentiation (LTP) and Ca^2+^ channels (Morgan and Teyler, 1999). The inhibition of Ca^2+^ channels by Al^3+^ could lead to a reduction in Ca^2+^ influx, resulting in reduced release of some neurotransmitters, which might explain impaired LTP (Platt et al., 1995).

**Fig. 8.**
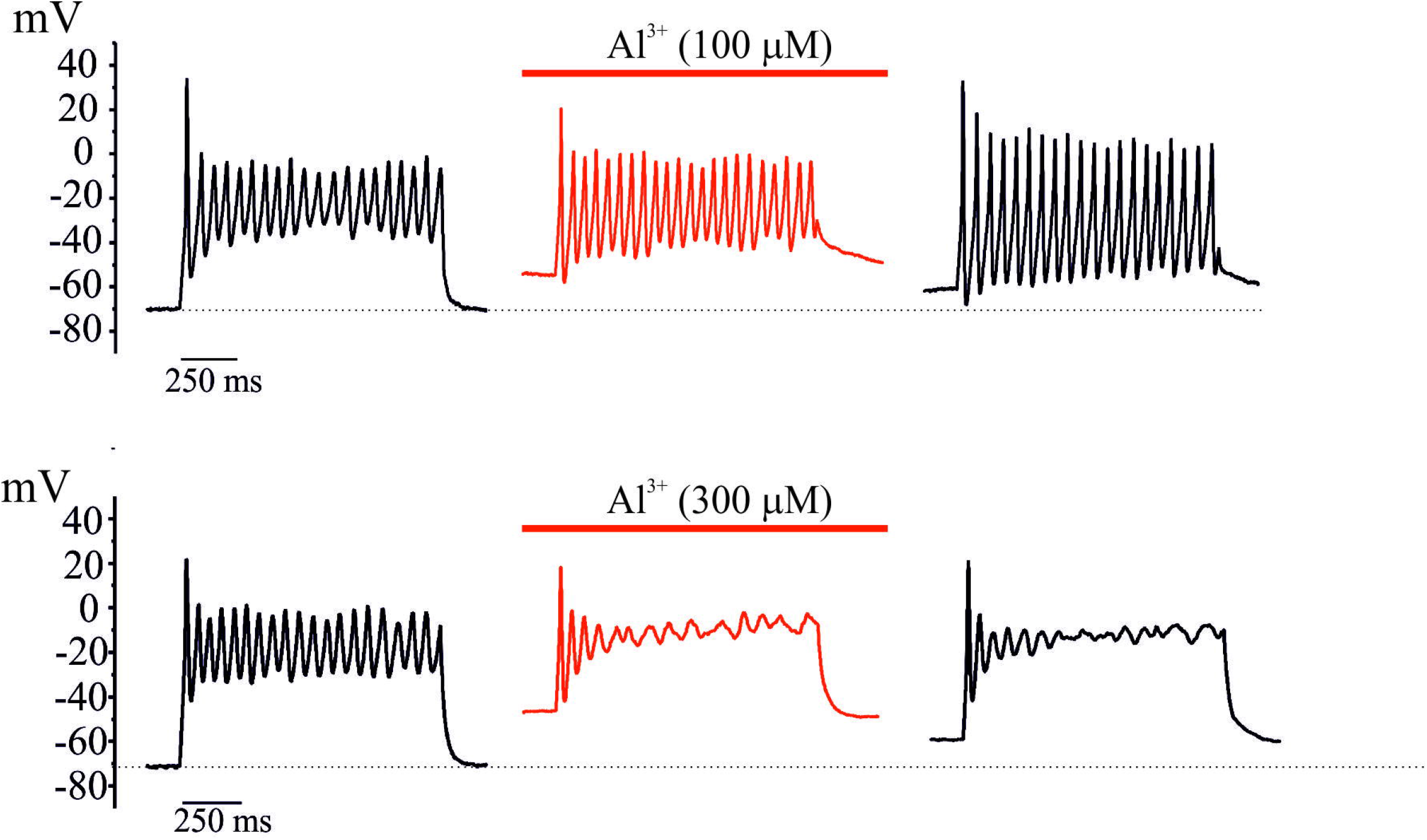
Depression of evoked APs exerted by Al^3+^. Current clamp recordings of evoked firing APs in response to low depolarizing injection currents (10 pA) applied during 1000 ms in control conditions and after Al^3+^ administration (100 and 300 μM).

## 5. Conclusion

In summary, we have evaluated the effects of acute aluminium application on calcium homeostasis and ionic currents, as responsible for neurosecretion and cellular excitability. In bovine chromaffin cells, Al^3+^ produced (1) an irreversible inhibition of catecholamine release, (2) a reduction of intracellular calcium, (3) a concentration-dependent and irreversible blockade of calcium currents, (4) a concentration-dependent and reversible blockade of sodium currents, (5) no significant effect on voltage-dependent potassium currents, (6) a concentration-dependent and irreversible blockade of calcium/voltage-dependent potassium currents, and finally, (7) an depression of the action potentials firing. Taking these results into consideration, Al^3+^ acts as a neurotoxic compound, affecting the cellular excitability and synaptic transmission.

## Abbreviations

AHP: afterhyperpolarization;
APs: action potentials;
BK: voltage and Ca^2+^-activated K^+^-channels;
CNS: central nervous system;
GABA: γ-aminobutyric acid;
I/V curve: intensity/voltage curve;
KRB: Krebs–Ringer bicarbonate;
NIFE: nifedipine;
RCCs: rat adrenal chromaffin cells;
SK: small conductance K^+^-channels;
TEA: tetraethylammonium;
TTX: tetrodotoxin;
ω: Aga-IVA,
ω: agatoxin-IVA;
ω: CTx-GVIA,
ω: conotoxin-GVIA;
ZnT3: Zn^2+^-transporter 3.

## Acknowledgments, funding sources and conflict of interest disclosure

This work was supported by MICIU (grant number PID2021-128133NB-I00/AEI/FEDER10.13039/501100011033). We also thank Fundación Instituto Teófilo Hernando for continued support. The authors declare that they have any financial interests in relation with the present work.

